# Spatiotemporal mapping of gene expression landscapes and developmental trajectories during zebrafish embryogenesis

**DOI:** 10.1101/2021.10.21.465298

**Authors:** Chang Liu, Rui Li, Young Li, Xiumei Lin, Shuowen Wang, Qun Liu, Kaichen Zhao, Xueqian Yang, Xuyang Shi, Yuting Ma, Chenyu Pei, Hui Wang, Wendai Bao, Junhou Hui, Michael Arman Berberoglu, Sunil Kumar Sahu, Miguel A. Esteban, Kailong Ma, Guangyi Fan, Yuxiang Li, Shiping Liu, Ao Chen, Xun Xu, Zhiqiang Dong, Longqi Liu

**Affiliations:** BGI-Shenzhen, Shenzhen 518083, China; College of Biomedicine and Health, College of Life Science and Technology, Huazhong Agricultural University, Wuhan, Hubei, 430070, China; Brain Research Institute, Taihe Hospital, Hubei University of Medicine, Shiyan, Hubei, 442000, China; College of Life Sciences, University of Chinese Academy of Sciences, Beijing 100049, China; BGI-Qingdao, BGI-Shenzhen, Qingdao, 266555, China; Department of Biology, University of Copenhagen, Copenhagen DK-2200, Denmark; Shenzhen Key Laboratory of Single-Cell Omics, Shenzhen 518083, China; Laboratory of Integrative Biology, Guangzhou Institutes of Biomedicine and Health, Chinese Academy of Sciences, Guangzhou 510530, China; CAS Key Laboratory of Regenerative Biology and Guangdong Provincial Key Laboratory of Stem Cells and Regenerative Medicine, Guangzhou Institutes of Biomedicine and Health, Guangzhou 510530, China; Institute of Stem Cells and Regeneration, Chinese Academy of Sciences, Beijing 100101, China

**Author notes:** Correspondence and requests for materials should be addressed to X.X., Z.D. or L.L. These authors contributed equally to this paper (Chang Liu, Rui Li, Young Li, Xiumei Lin, Shuowen Wang, Qun Liu).

**Keywords:** Stereo-seq, spatial transcriptomics, scRNA-seq, embryonic development, zebrafish

## Abstract

Vertebrate embryogenesis is a remarkably dynamic process during which numerous cell types of different lineages generate, change, or disappear within a short period of time. A major challenge in understanding this process is the lack of topographical transcriptomic information that can help correlate microenvironmental cues within the hierarchy of cell fate decisions. Here, we employed Stereo-seq, a high-definition spatially resolved transcriptomic technology, to dissect the spatiotemporal dynamics of gene expression and regulatory networks in the developing zebrafish embryos. We profiled 91 embryo sections covering six critical time points during the first 24 hours of development, obtaining a total of 139,391 spots at cellular size (∼100 μm^2^) with spatial coordinates. Meanwhile, we identified spatial modules and co-varying genes for specific tissue organizations. By performing the integrative analysis of the Stereo-seq and scRNA-seq data from each time point, we reconstructed the spatially resolved developmental trajectories of cell fate transitions and molecular changes during zebrafish embryogenesis. We further investigated the spatial distribution of ligand-receptor pairs for major signaling pathways and identified novel interactions that potentially crosstalk with the Notch signaling pathway during zebrafish development. Our study constitutes a fundamental reference for further studies aiming to understand vertebrate development.

## INTRODUCTION

Vertebrate embryogenesis is an intricate and dynamic process with intense gene expression changes and frequent cell state transitions within short time windows. Extrinsic and intrinsic cues, including transcription factors (TFs), morphogens, signaling pathways, and signals from the extracellular matrix (ECM), play pivotal roles in determining different cell fates which present distinct morphologies, spatial positions and functions (Bardot and Hadjantonakis, 2020; Marlow, 2020; Vining and Mooney, 2017). How these regulatory factors spatially interact and function together to induce a complex vertebrate embryo in a precisely controlled manner is one of the fundamental questions about embryogenesis that demands further investigation. In particular, understanding how ligand-receptor pairs spatially interact to switch on/off specific signaling pathways, such as Notch, Wnt, SHH and TGF-b during vertebrate embryogenesis is crucial but poorly documented.

The zebrafish is a widely-used model organism for studying vertebrate embryonic development thanks to its fast development, embryonic transparency, and accessibility to both physical and genetic manipulation. The advances in sequencing technologies have made possible the assembly of single-cell atlases of model organisms during development, which enables genome-wide profiling of multimodal information including gene expression, epigenetic state, and protein level in individual cells (Cao et al., 2017; Han et al., 2020; Karaiskos et al., 2017; Stelzer et al., 2015; The Tabula Muris Consortium et al., 2018; Trevino et al., 2020). Efforts have been made to map the gene expression landscapes during zebrafish embryogenesis and developmental trajectories have been constructed, which defined the transcriptomic states of cells as they acquire their fates (Briggs et al., 2018; Farrell et al., 2018; Wagner et al., 2018). However, how the time-course transcriptomic states correlate with each other in the background of spatial localization in a complex developing zebrafish embryo remains elusive and the spatial organization of different cell types in complex tissues remains poorly understood due to the limitation of current spatial transcriptomic technologies (Li et al., 2021; Liu et al., 2020; Rodriques et al., 2020; Stickels et al., 2021; Vickovic et al., 2019).

Based on the traditional *in situ* hybridization technology, a representative database of gene expression in the developing zebrafish embryos and larvae (The Zebrafish Information Network (ZFIN)) (Sprague, 2003) is available for scientists to inquire the expression pattern of a specific gene, but the *in situ* images lack cell type information and the global changes of transcripts cannot be explored. A spatial transcriptomic technology named Tomo-seq has been applied in zebrafish embryos, but this technology is based on bulk sequencing of cryosections and the spatial information on the tissue sections is missing (Holler et al., 2021; Junker et al., 2014). Here, we adopted the newly developed SpaTial Enhanced REsolution Omics-sequencing (Stereo-seq) (Chen et al., 2021) to dissect the spatiotemporal gene expression landscapes with high resolution and sensitivity during the first day of zebrafish embryo development. Our study provides important data resources for the research of gene expression, cellular organization and regulatory networks during zebrafish embryogenesis.

## RESULTS

### Generation of high-resolution spatiotemporal transcriptomic data by Stereo-seq in zebrafish embryos

Zebrafish embryos of different stages (3.3 hours post-fertilization (hpf), 5.25 hpf, 10 hpf, 12 hpf, 18 hpf, 24 hpf) were harvested for spatial transcriptomic analysis (Figure 1A, top). Embryos were dechorionated and embedded in optimal cutting temperature (OCT) compound for the preparation of sagittal cryosections with a thickness of 15 μm (close to cellular size). High resolution Stereo-seq chips (spot size 220 nm, center-to-center size 715 nm, chip size 1 cm^2^) were used for the capture of RNA transcripts (Figure 1A, bottom). In order to avoid incomplete sampling of cell types and batch effects, we obtained multiple sections of one embryo at each stage for Stereo-seq. Taking 24 hpf as an example, 17 sections of the same embryo were attached on two 1 cm^2^ chips and subjected to Stereo-seq (Figure 1B-1C). We applied a grid-based strategy, which aggregates a number of DNA nanoball (DNB) spots, to cluster the spatial signature into substructures (see Methods). As expected, by comparison of spatial clustering with different grid (bin) size ranging from 143 μm (bin 200, 200 × 200 DNB) to 10 μm (bin 15, 15 × 15 DNB, equivalent to ∼1 cell diameter), we found more refined clustering with increased cluster number and accuracy was achieved by employing a higher-resolution bin (Figure 1D). Meanwhile, the number of transcripts and genes detected among different sections of the 24 hpf embryo were comparable (Figure 1E-1F), and we recovered a median of 1250 unique molecular identifiers (UMIs) and 394 genes with bin 15 (Figure S1A). The number of embryo sections and the average numbers of UMIs and genes were overall sufficient for further analysis across all the 6 developmental stages (Figure 1G and S1A).

**Figure 1.**
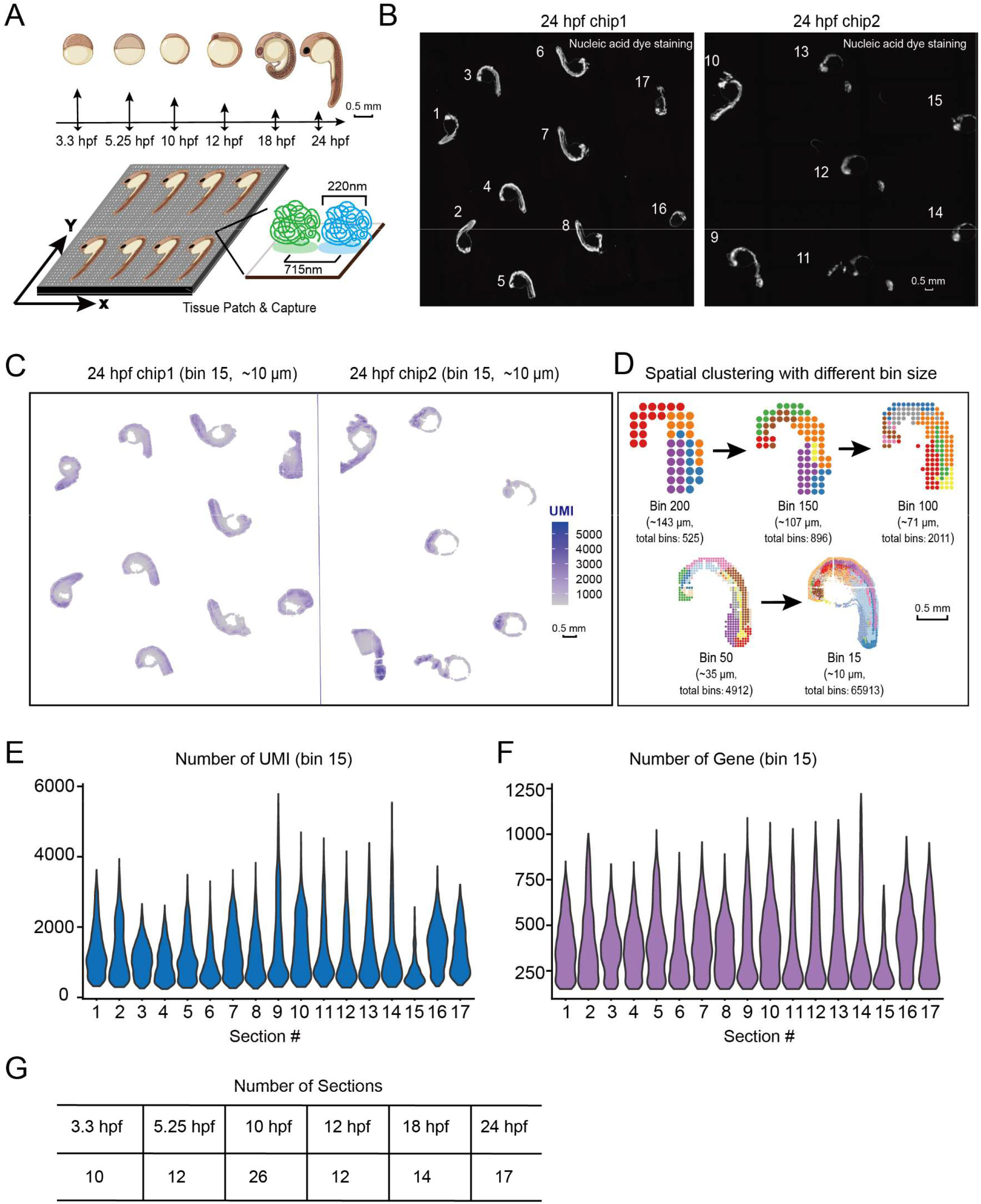
High resolution Stereo-seq on multiple sections of the developing zebrafish embryos. (A) Experimental outline (top): diagram of the zebrafish embryos at different development stages which were subjected to Stereo-seq. Scale bar: 0.5 mm. Stereo-seq process diagram (bottom): the enlarged image shows the size of each spot and the distance between 2 adjacent spots. (B) Nucleic acid dye staining of the 24 hpf zebrafish embryo sections attached on two 1-square-centimeter Stereo-seq chips. Scale bar: 0.5 mm. (C) Spatial visualization of the distribution of captured transcripts (unique molecular identifiers (UMIs)) on all 24 hpf zebrafish embryo sections. Scale bar: 0.5 mm. (D) Unsupervised clustering of the 24 hpf zebrafish embryo section analyzed by Stereo-seq at different bin sizes. Scale bar: 0.5 mm. (E and F) Violin plot of the number of captured transcripts (E) and genes (F) of each 24hpf zebrafish embryo sections. (G) Table summarizing the number of sections used in Stereo-seq at each embryonic development stage.

Thus, our Stereo-seq with multiple-section strategy enables high-resolution and comprehensive spatial transcriptomic analysis of small-sized samples such as zebrafish embryos.

### Spatial clustering and molecular characterization of the zebrafish embryogenesis spatiotemporal transcriptomic atlas

Stereo-seq with multiple-section strategy at cellular size resolution (bin 15) generated high-definition spatial transcriptomic data from developing zebrafish embryos, which can hardly be profiled by other available technologies due to a much lower density of signatures. This allowed us to create a high-quality Zebrafish Embryogenesis Spatiotemporal Transcriptomic Atlas (ZESTA).

We pooled all the sections of each stage for unsupervised clustering (see Methods), and revealed spatial heterogeneity of the embryos and spatial specification dynamics of different regions during development (Figure 2A). As Stereo-seq captured a high density of signals, we were able to identify more elaborate structures with deeper clustering (Figure 2B and S1B-S1G). Some structures can be further divided into detailed substructures. For example, the brain components from the Stereo-seq data of the 24 hpf embryo could be further divided into subdivisions such as ‘telencephalon’ and ‘dorsal diencephalon’ (Figure 2C). Similarly, the eye can be divided into ‘optic lens’ and ‘pigment-epithelium’ (Figure 2C). On the other hand, the spinal cord, floor plate, and notochord are adjacent interacting structures and could be separated precisely on the embryo sections by specific expression of genes including *prdm8, ntn1b* and *ntd5*, respectively (Figure 2C). Gene expression kinetics across all stages revealed restricted expression at specific time points and anatomic regions, as exemplified by erythroid lineage cell (*hbbe1*.*1*), myotome (*tnnc2*), yolk syncytial layer (YSL) (*apoa1a*) and hatching gland (*he1*.*1*) (Figure 2D). Genes of early embryogenesis, i.e., *pou5f3* and *s100a1* showed specific expression at early stages, consistent with known expression patterns identified by classic *in situ* hybridization experiments (Figure 2D) (Diks et al., 2006; Onichtchouk and Driever, 2016).

**Figure 2.**
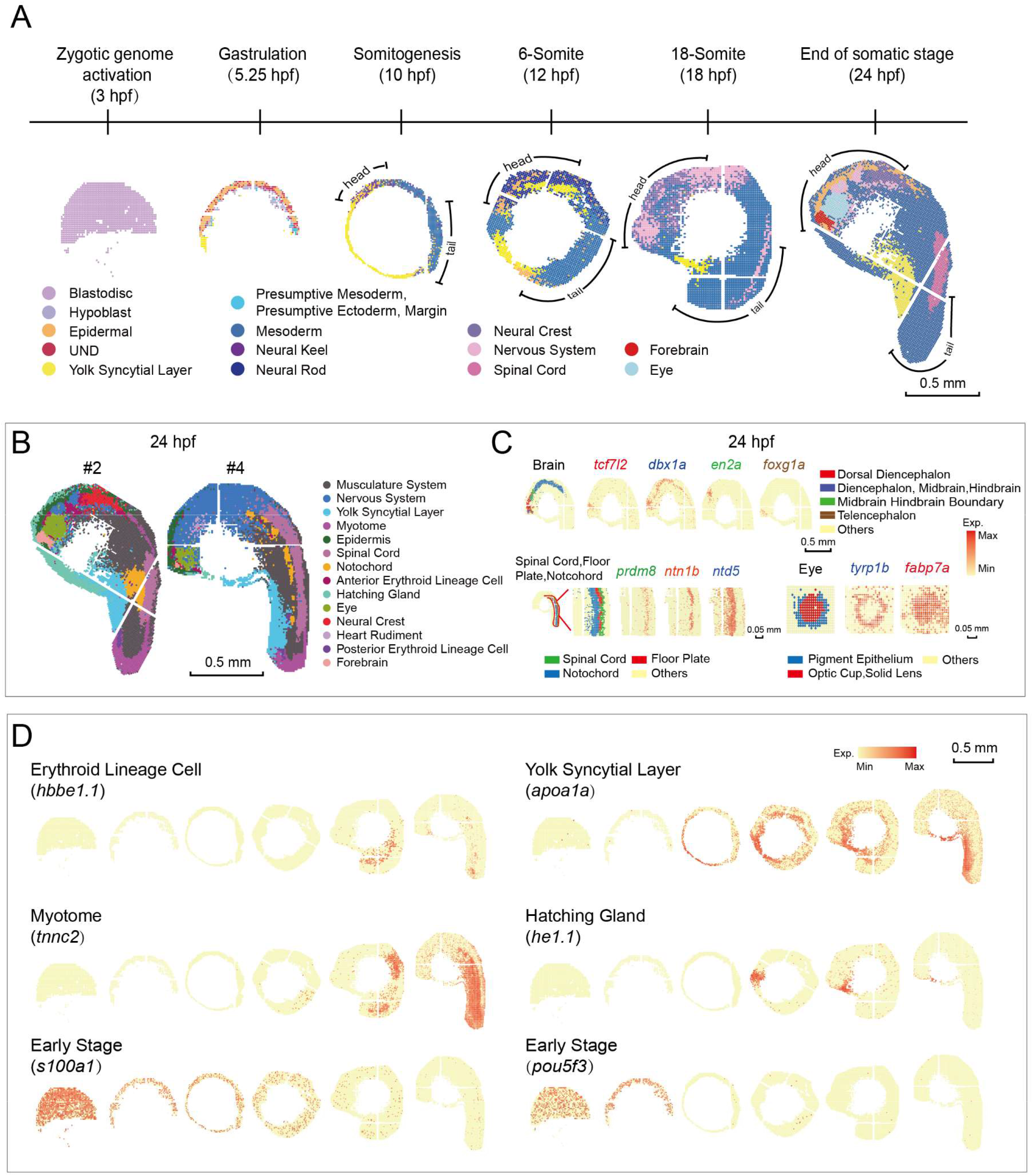
A spatial transcriptomic atlas at cellular size resolution of the developing zebrafish embryo. (A) Unsupervised clustering of the zebrafish embryo section across sequential developmental stages analyzed by Stereo-seq at bin 15. Cells are colored by different regions. (B) Unsupervised clustering of the 24hpf zebrafish embryo sections at bin 15. Cells are colored by spatial identities inferred from expressed marker genes. Scale bars 0.5 mm. (C) Spatial visualization of indicated areas of the 24 hpf embryo on the left: detailed anatomical structures identified in the brain and the eye, and a combined structure including the spinal cord, the floor plate, and the notochord. The expression patterns of marker genes for each anatomical structure are shown on the right. Scale bars: 0.5 mm and 0.05 mm. (D) Spatial visualization of the expression of indicated genes for erythroid lineage cell, myotome, YSL, hatching gland and early development stages. Scale bar: 0.5 mm.

Taken together, our Stereo-seq constituted a unique resource of ZESTA with unprecedented resolution, and the publicly available interactive data portal can be accessed at http://stereomap.cngb.org/zebrafish/data_index.

### Spatial gene modules uncover the interaction of different spatial regions

The spatial patterns of gene expression depict different functional regions across embryo sections. In order to identify co-varying genes showing similar spatial distribution, we applied Hotspot (DeTomaso and Yosef, 2021) to identify gene modules on our Stereo-seq dataset. We identified 12 spatial modules for the 24 hpf embryo sections. Gene Ontology (GO) enrichment analysis on the spatially correlated genes of each module confirmed the functions of these spatial modules (Figure 3A), which also showed consistency with the spatial clusters (Figure 3B). Spatial visualization of modules showed that the distribution of spatial modules reasonably matched their region-specific biological functions (Figure 3A and 3C). For example, module 7 (M7, posterior erythroid lineage cell) gathered genes involved in oxygen transport, and M8 (eye) was abundant in genes related to lens development. For the other developmental stages, we identified 8 spatial modules for 10 hpf, 11 for 12 hpf and 16 for 18 hpf (Figure S2A, S2D and S2G). Similarly, the GO function enrichment, spatial location and cell types reasonably matched with each other for most spatial modules (Figure S2B-C, S2E-F and S2H-I).

**Figure 3.**
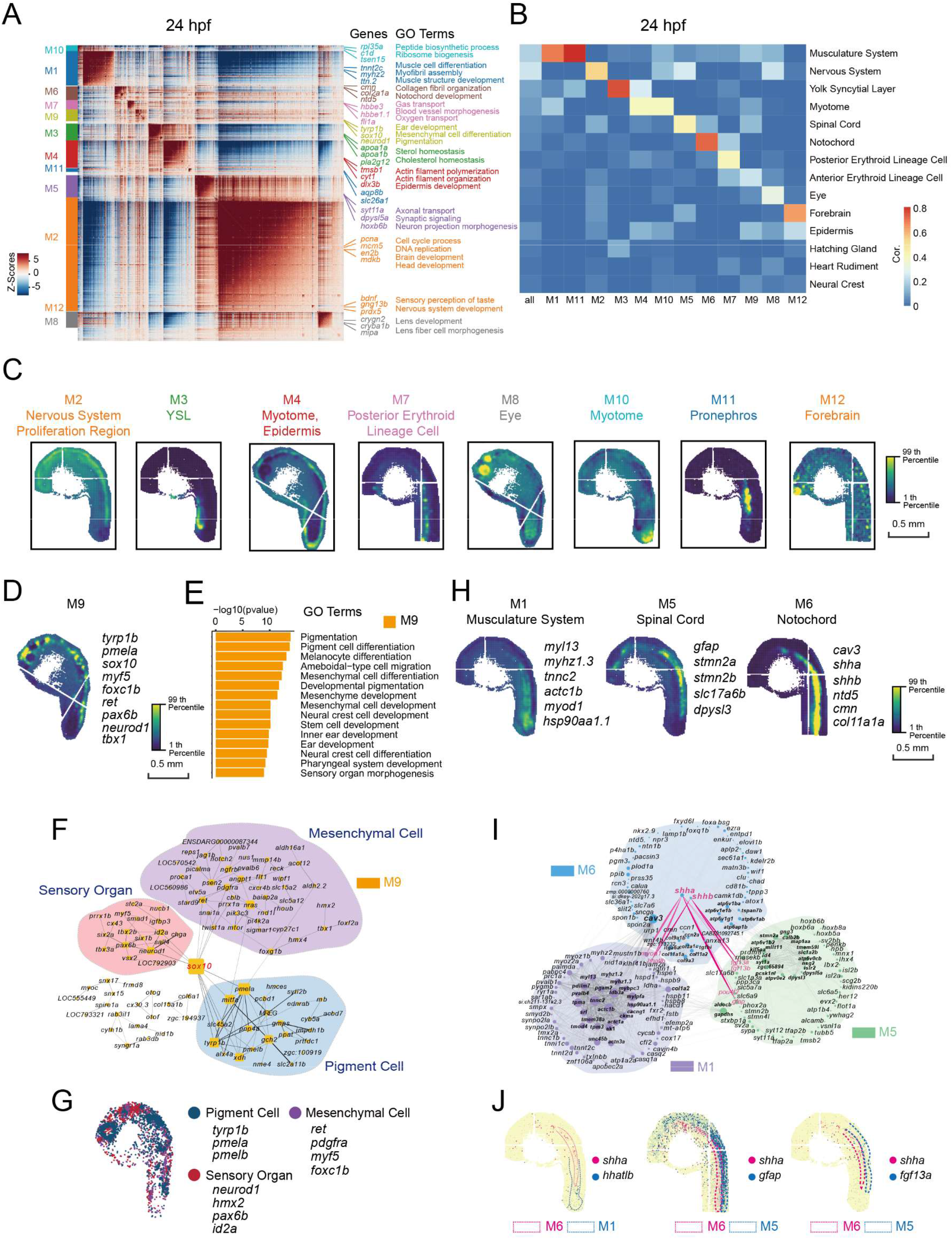
Spatial modules identified by Hotspot uncover the interaction among spatial regions in 24 hpf zebrafish embryo. (A) Heatmap shows the genes with significant spatial autocorrelation (5,089 genes, FDR<0.05) grouped into 12 gene modules based on pairwise spatial correlations of gene expression in multiple sections of the 24 hpf zebrafish embryo. Selected genes and GO term related to representative modules are highlight on the right side. (B) Heatmap shows the Pearson correlation of the module score for each spatial autocorrelation module and the expression sets of specific genes for each spatial cluster of the 24 hpf embryo from Stereo-seq dataset in Figure 2B. (C) Spatial visualization of representative modules on 24 hpf embryo sections. Scale bars 0.5 mm. (D) Spatial visualization of module 9 (M9). (E) Bar graph shows significantly enriched selected gene ontology terms in M9. (F) Protein-protein interaction (PPI) network of genes in M9. The network is visualized by Cytoscape. Node size represents the relative connectivity, and the thickness of the dark line represents the strength of interactions. (G) Spatial expression pattern of genes in M9. These genes are related to the development of pigment cell (blue), mesenchymal cell (purple), and sensory organ (red). (H) Spatial visualization of modules related to notochord and neighboring tissues on 24 hpf embryo sections including M1, M5 and M6. (I) PPI network of genes in the 3 modules in H. The red lines represent interactions with *shha/b*. (J) Spatial expression patterns of *shha* and the interactive genes.

Interestingly, we found that a single gene module can correspond to multiple subregions. For instance, M9 in 24 hpf embryo spread across various clusters including neural crest, erythroid lineage cell, epidermis, and eye (Figure 3B and 3D). GO enrichment of this module revealed functions related to the development of pigment cells, mesenchymal cells, and sensory organ (Figure 3E). In order to explore the connection among these three cell types/organs, STRING was employed to seek potential gene interaction which was visualized with Cytoscape (Figure 3F). Intriguingly, we found that the transcription factor gene *sox10* intensively interacts with genes from these three cell types/organs, consistent with the reported function of *sox10* (Rocha et al., 2020). As expected, projection of specific genes from pigment cell, sensory organ, and mesenchymal cell on the same embryo section revealed spatial neighboring distributions (Figure 3G).

The notochord provides directional signals to the surrounding tissues during development through secreting sonic hedgehog (SHH), a key morphogen regulating organogenesis (Corallo et al., 2015; Male et al., 2020). To investigate the interaction of notochord with the neighboring regions, we constructed the interaction network of genes in M6 (notochord (Figure 3A and 3H, right)) and the neighboring modules, M1 and M5 which correspond to the musculature system and the spinal cord respectively (Figure 3A and 3H, left). Within the potential interacting genes revealed by STRING, we found that *myod1* and *hhatlb* in M1 showed interaction with *shha* and *shhb* in M6. Previous studies have found that *shh* plays a role in muscle development and adult myogenesis by regulating the activity of *myod1* (Voronova et al., 2013), while to the best of our knowledge, *hhatlb* has been rarely reported to interact with *shh*. On the other hand, *fgf13* and two other neural stem cells maker genes (*gfap* and *pou4f2*) in M5 show interaction with *shha/b*. The interaction of these genes was further confirmed by their tightly adjacent characters of spatial distribution (Figure 3J).

In summary, spatial modules depict the functional regions or subregions in an organism, provide a powerful tool to explore the genetic interaction between different cell types, and serve as a reliable resource to discover novel function as well as unknown interactions of genes.

### Construction of spatially resolved developmental trajectories by integrated analysis of scRNA-seq and Stereo-seq data

Both our Stereo-seq data and previously published scRNA-seq data indicate that developmental time is a strong source of variation mainly due to active transition between different cell states during embryogenesis. In order to construct the spatiotemporal developmental trajectories of embryonic cells, we further performed droplet based scRNA-seq (Liu et al., 2019) (see Methods) with zebrafish embryos at the same developmental time points as in the Stereo-seq profiling (Figure 4A). We obtained 86,307 cells from embryos of the 6 developmental stages and identified 71 cell types, which are mostly consistent with previous studies (Farrell et al., 2018; Wagner et al., 2018) (Figure 4A and S3A-S3H).

**Figure 4.**
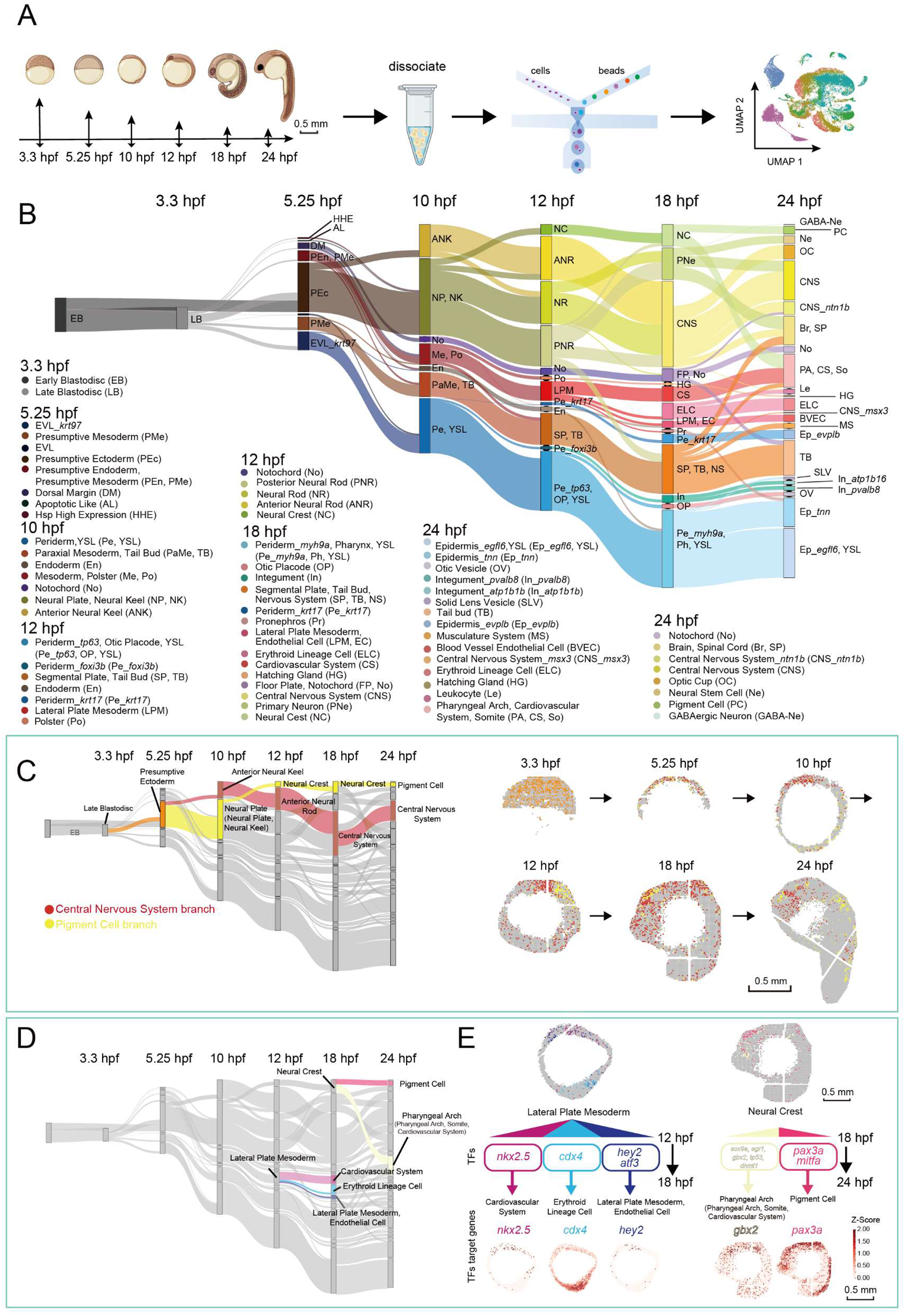
Construction of the spatially resolved developmental trajectories by integrated analysis of the scRNA-seq and Stereo-seq data. (A) Schematic representation of the workflow for scRNA-seq of zebrafish embryos at different developmental stages using the C4 system. (B) Application of Sankey diagram to visualize zebrafish embryo developing trajectory with scRNA-seq data. The abbreviations of the cell types in B are also marked in parentheses after each cell type in the color legend. (C) Application of SPOTlight to integrate Stereo-seq with scRNA-seq data to infer the spatially resolved developmental trajectories. Two developmental branches which are developed from the presumptive ectoderm, namely, pigment cell, and central nervous system (left) are simultaneously displayed on six sequential spatial sections to show the different spatial developmental trajectories (right). (D) Two selected developmental branching points at 12 hpf and 18 hpf. The lateral plate mesoderm (left) and neural crest (right), are shown in the Sankey diagram. (E) Cell fate regulatory maps of different developmental destinations at 12 hpf and 18 hpf related to Figure 4D. SPOTlight was applied to visualize the spatial distribution of cell subgroups with different differentiation (top); selected representative crucial TFs for different developmental branches are shown in the chart (middle); the spatial expression distributions of TFs target genes are visualized on embryonic sections (bottom). The scale bars represent 0.5 mm.

At 3.3 hpf, the emergence of two cell types was observed, for which we used Monocle 2 to undertake pseudo-chronological analysis to determine their differential identities. Based on the pseudo-chronological relationship, we determined the two cell types as early blastodisc and late blastodisc respectively (Figure S4B). To construct the developmental trajectory, we integrated scRNA-seq data of adjacent time points using Harmony (Korsunsky et al., 2019) and applied KNN mapping to predict the developmental fate of each cell of the earlier time point (see Methods). We then projected the developing trajectory using the Sankey diagram (Figure 4B).

Although single-cell studies have mapped the cell types and their molecular features during vertebrate embryogenesis, the dynamics of the spatial organization of cells is still poorly understood (Farrell et al., 2018; Wagner et al., 2018). To introduce spatial information to the cell fate lineages, we conducted an integrative analysis with the combination of the scRNA-seq and Stereo-seq data by applying SPOTlight (Elosua-Bayes et al., 2021) to calculate the cell composition of each bin in the Stereo-seq data, which allowed projection of the developmental trajectories to the embryo sections (Figure 4C and S4A-S4B). These results revealed the spatial dynamics of different developmental trajectories. Taking the two developmental branches of the presumptive ectoderm as an example, the central nervous system branch and the pigment cell branch both developed from the presumptive ectoderm which appeared at 5.25 hpf with a discrete distribution pattern, and the two branches started to be distinguished at 10 hpf. The central nervous system branch showed an anteriorly concentrated pattern starting at 10 hpf and enriched in the brain region with some leakage at the spinal cord. However, the pigment cell branch remained discrete on the section at 10 hpf while appeared to gather at the anterior part of the embryo section at 12 hpf when the neural crest cell fate was evident and subsequently differentiated into pigment cells (Figure 4C and S4A).

A prominent phenomenon of the developmental trajectory is that some cell types bifurcate into multiple branches at certain time points (Figure 4D and S4C). We noticed that the distributions of some developmental directions seem to be already spatially separated and distinguishable in the progenitor cell type (Figure 4E, top and S4D, top). We integrated the single-cell transcriptomic data and applied SCENIC (Aibar et al., 2017) algorithm to predict TFs with high activity at each developmental directions (see Methods) (Figure 4E, middle and S4D, middle). For example, different TFs were identified to regulate the segregated developmental directions of the 12 hpf lateral plate mesoderm, which gives rise to the cardiovascular system, erythroid lineage cell and endothelial cell. The *nkx2*.*5* gene is predicted as crucial TFs for the development of the cardiovascular system, while *cdx4* for erythroid lineage cell (Figure 4E, left). Similarly, different TFs were identified for the separate developmental lineages of the 18 hpf neural crest (Figure 4E, right). We visualized the regulatory activities of these key TFs by displaying the target gene expression on embryo sections, for which the distribution of target genes is largely consistent with that of the corresponding cell type (Figure 4E, bottom and S4D, bottom).

As described above, our spatiotemporal developmental trajectories revealed the dynamic cell fate transitions with anatomical annotation during zebrafish early embryogenesis, and can be used to investigate potential regulatory TFs for a specific developmental direction.

### Dynamic spatiotemporal distribution of ligand-receptor pairs during zebrafish embryogenesis

The extrinsic cues, including the interaction between ligands and receptors, are crucial for the regulation of cell fate during embryogenesis. We, therefore, calculated the relative distance of different ligand-receptor pairs at all six developmental stages with the distance of adjacent bins (Figure 5A and S5A-S5B). Considering the dropout in the transcriptomic data, we define that the ligand and receptor have strong potential to interact when their average relative distance was less than 5 bins at least at one time point. Combining the relative distance and expression strength, we illustrated the dynamics of the ligand-receptor interactions during zebrafish embryonic development (Figure 5B). We observed an increasing number of receptor-ligand pairs that come to interact as the embryo develops. Moreover, the expressions of most active ligand-receptor pairs were upregulated and the distances became smaller.

**Figure 5.**
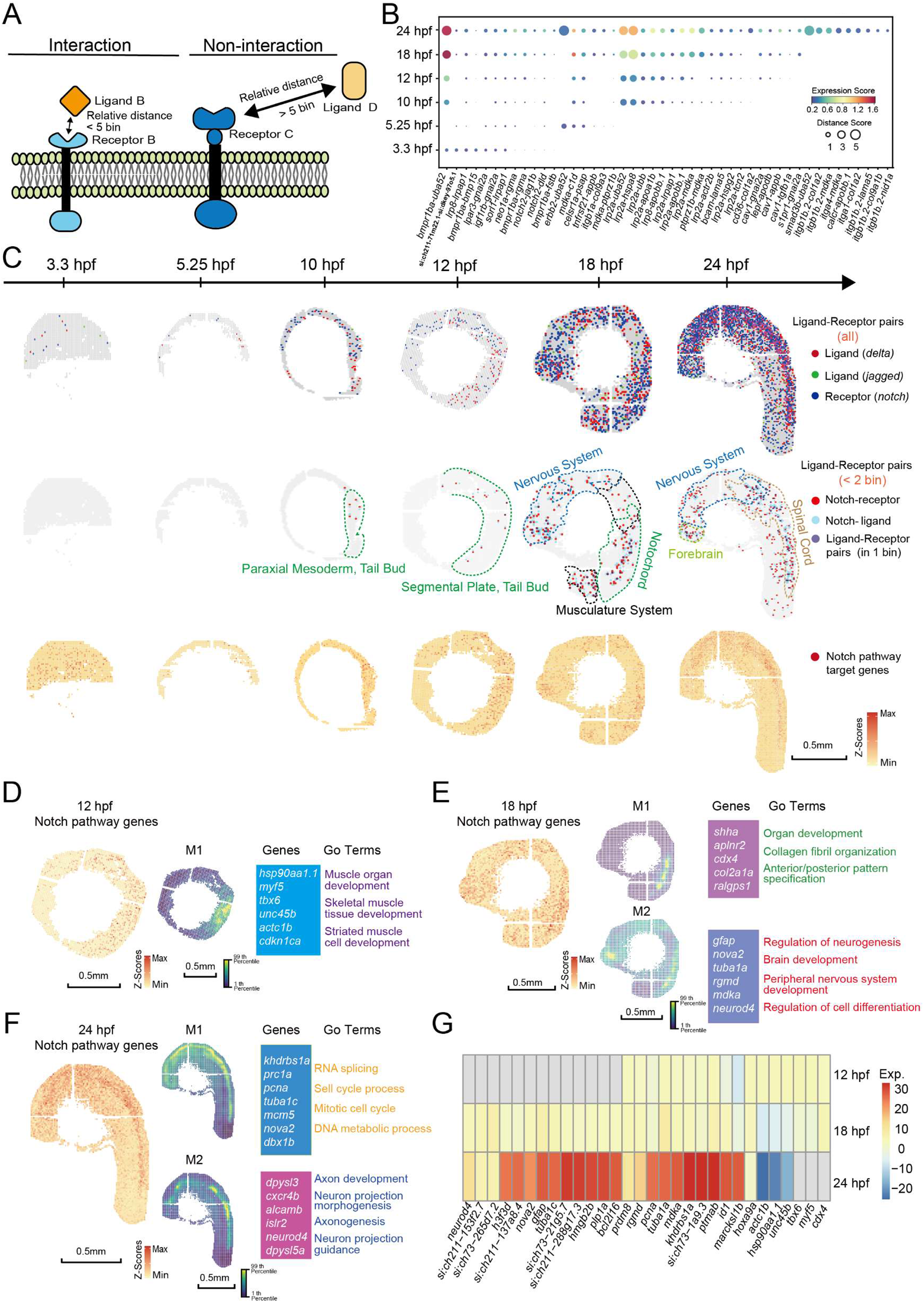
The relative spatial distance of different ligand-receptor pairs reveals the arrangement and interaction of signaling pathways. (A) Model diagram of the analysis of ligand-receptor relative distance. (B) A dotted heat map shows the expression score and relative distance score of ligand-receptor pairs. (C) Spatial expression pattern of *notch* family genes (blue), *deltal* family genes (red), and *jagged* family genes (green) on embryo sections at the 6 time points (top). Spatial expression pattern of Notch receptors (red), ligands (blue) with a relative distance less than 2, and the ligand-receptor pairs expressed in the same bin (purple) (middle). Spatial expression pattern of Notch target genes (red) on the embryonic sections at different time points (bottom). Scale bar: 0.5 mm. (D-F) Gene modules identified using Hotspot that showed spatial correlation with Notch pathway in 12 hpf (D), 18 hpf (E), and 24 hpf (F) embryos. The spatial expression pattern of Notch pathway genes at each developmental stage is shown on the left, and selected genes and GO terms related to spatially correlated modules are highlight on the right. Scale bars: 0.5 mm. (G) A heatmap shows the temporal expression change of genes selected from modules in D-F.

The Notch signaling pathway is conserved in most vertebrates and plays an essential role in cell fate determination through lateral inhibition (Andersson et al., 2011). We graphed the distance heat map of the Notch ligand-receptor pairs at different developmental stages and found that multiple pairs showed time-dependent increase in proximity (Figure S5C). Then we explored the dynamic spatial distribution of the Notch components at all six time points, which revealed that the Notch receptors and ligands (*delta, jagged*) were sparsely expressed at 3.3 hpf and 5.25 hpf, and the expression was increased at 10 hpf without any obvious pattern. Higher Notch ligand-receptor proximity was observed at the somite forming site on 12 hpf sections. The expression of both ligands and receptors showed wider distribution but was enriched in the brain and the somite at 18 hpf, and was mainly concentrated in the nervous system at 24 hpf (Figure 5C, top). To illustrate the spatial ligand-receptor interactions of Notch signaling, we selected the receptor-expressing bins with a relative distance less than 2 bins to the ligands, plotted them together with the nearest ligand-expressing bins on the embryo sections (Figure 5C, middle), and illustrated the corresponding cell clusters (Figure S5D). We further analyzed Notch activities by mapping the expression of downstream target genes (Figure 5C, bottom), which is consistent with the spatial expression dynamics of Notch ligand-receptor pairs. Our results revealed a highly dynamic spatial distribution of Notch components expression and Notch activity during zebrafish embryogenesis.

We further explored gene modules that are spatially correlated with Notch signaling pathway using Hotspot analysis on the Stereo-seq data of 12 hpf, 18 hpf, and 24 hpf zebrafish embryos (Figure 5D-5F) (see Methods). At 12 hpf, one Notch correlated spatial module is identified with distribution in the somite-forming region and contains genes such as *myf5, unc45b* and *hsp90aa1*.*1* which are functionally associated with muscle development (Figure 5D). Two Notch correlated spatial modules showed up at 18 hpf (Figure 5E). M1 gathers in the somitogenesis region and is abundant in genes that function in organ development, collagen fibril organization and anterior-posterior pattern specification such as *shha, cdx4* and *col2a1a*. M2 is concentrated in the central nervous system comprising genes that are important in neurogenesis and brain development, *i*.*e. gfap* and *neurod4*, which regulate gliogenesis and neuronal differentiation respectively (Fukuoka et al., 2021) . At 24 hpf, the Notch correlated spatial modules are also enriched in the nervous system (Figure 5F). M1 contains genes such as *pcna, mcm5* and *nova2*, which are related to cell cycle, RNA splicing and DNA metabolism, indicating that the dominant role of this module is to regulate the development of neural progenitor cells (Ino and Chiba, 2000; Yano et al., 2010). In contrast, genes involved in M2 such as *dpysl3, cxcr4b*, mainly function in axonogenesis and neuronal projection development. Consistent with spatial expression dynamics of Notch ligand-receptor pairs, correlated spatial modules showed that the Notch correlated genes were weak and enriched at the somitogenesis region at 12 hpf, then distributed widely but were enriched in the brain and at the somitogenesis region at 18 hpf, and were concentrated in the nervous system at 24 hpf (Figure 5D-5F). In order to identify candidate genes that directly or indirectly interact with the Notch pathway at each developmental stage, we further characterized the co-varying tendency of module genes relative to the Notch pathway at 12 hpf, 18 hpf and 24 hpf (Figure 5G). Therefore, the spatial gene modules showed co-varying expression patterns and functional correlation with Notch signaling pathway at specific time points, which indicates that the genes involved can very likely interact with Notch signaling, either directly or indirectly, to play an important role in the development of defined cell types.

We thereby demonstrated a spatiotemporal expression atlas of various ligand-receptor pairs and provided a useful database for the investigation of the dynamics of different signaling pathways as exemplified by the comprehensive analysis on Notch pathway.

## DISCUSSION

The mapping of a spatial transcriptomic landscape at cellular resolution is essential for understanding vertebrate embryogenesis. Due to the limitation of current spatial transcriptomic technologies, there is a lack of high-resolution spatial transcriptomic resources for the zebrafish embryos, a popular model organism for developmental biology. The newly developed Stereo-seq enables the depiction of the spatial transcriptomic landscape at cellular resolution with high sensitivity during zebrafish embryogenesis (Chen et al., 2021). In addition, the flexible size of the Stereo-seq chip allows us to simultaneously attach multiple sections on one chip, which can greatly eliminate incomplete sampling of cell types and batch effects introduced by separate experimental runs. The present study, for the first time, employed the Stereo-seq, and drafted the dynamic spatiotemporal landscape of gene expression as well as spatial regulatory factors at single-cell size resolution during the first day of zebrafish embryonic development. Our high-resolution Stereo-seq data successfully distinguished detailed anatomical structures such as the hypoblast, EVL, hatching gland, and spinal cord at different developmental stages and identified organ subdivisions such as the telencephalon, midbrain-hindbrain boundary, and dorsal diencephalon of the brain. Besides, certain structures and gene expression patterns can be further zoomed in by extracting the tissue-specific data from our online resource to perform personalized analyses (http://stereomap.cngb.org/zebrafish/data_index).

Taking advantage of our Stereo-seq data, we identified spatial modules at different developmental stages and integrated the spatial modules with spatial regions to investigate key co-varying genes. Nevertheless, how the spatially correlated gene sets interact and mutually regulate each other both inside and outside the modules to form a regulatory network remains elusive, and unraveling it would improve our understanding of the molecular mechanisms of vertebrate embryogenesis. For example, the notochord provides directional signals to the surrounding tissue during embryogenesis, in which SHH plays a critical role. Consistent with the known knowledge, our analysis of the spatial modules showed that the notochord interacts with the adjacent spinal cord and muscle through *shh*. In addition to *shh*, we found that *cav3* in M6 could interact with the genes related to muscle contraction in M1. The Cavin protein (including Cav1/Cav3) has been reported as essential for the development of notochord and muscle (Lim et al., 2017). We, therefore, discovered the potential interacting genes of *cav3* through analyzing the interaction of spatial module genes, which will provide the rationale for further mechanistic investigations on the function of *cav3* during organogenesis (Figure 3I).

To accurately map the cell type determination and developmental trajectory reconstruction, we have constructed the spatiotemporal developmental trajectories of the zebrafish embryo by combining the scRNA-seq and Stereo-seq data to delineate cell state transitions with spatial coordinates. The developmental trajectories showed clear and specific spatiotemporal characteristics, which are consistent with previously known facts about zebrafish embryonic development. Besides illuminating the spatial separation of closely correlated cell types, our spatiotemporal developmental trajectories resolved some previously unanswered questions. For instance, it has been unclear when the lateral plate mesoderm starts to differentiate into different tissues. Our results disclosed that the fate of this structure is already decided to differentiate in three different directions, namely, the cardiovascular system, erythroid lineage cell, and endothelial cell at 12 hpf (Figure 4E and S4A).

TFs, together with their downstream gene networks, are one of the key factors that drive cell fate transition during embryogenesis. We investigated candidate TFs that potentially dominate the cell types at different time points in our spatiotemporal developmental trajectories. Our data identified known TFs, whose spatiotemporal distribution was consistent with their reported functions. The TF gene *cdx4* has been shown to be involved in the development of hematopoietic stem cells (Davidson and Zon, 2006) and our analysis showed that *cdx4* regulates the cell fate transition from lateral plate mesoderm to erythroid lineage cell (Figure 4E, left). We also discovered a potential new role of certain TFs such as *foxn4* in the differentiation of the presumptive ectoderm from the blastodisc, while *foxn4* was previously known to function in the regulation of neural stem cell division (Misra et al., 2014) (Figure S4D).

Based on our high-resolution Stereo-seq data, we were able to precisely calculate the spatial distance between different ligands and receptors embryo-wide in zebrafish. Besides the Notch components, we uncovered interesting dynamics of many other ligand-receptor pairs. The ligand *uba52* has a strong interaction with multiple receptors at different developmental stages, e.g., interacting with the BMP receptor *bmpr1ba* from as early as 3.3 hpf while starting to interact with *lrp2a* at 10 hpf. The bmp family receptor (*bmpr1ba*) shows spatial proximity with the ligand as early as 3.3 hpf in our analysis, while the receptor gene *lrp2a* and the corresponding ligand genes are co-expressed at 10 hpf when the somite and the nervous system start to develop. The integrin gene *itgb1b*.*2*, which encodes the molecules that function to attach the cells to the ECM and transduce signals from the ECM to the cells, starts to get close to the ligands at 18-24 hpf. The highly dynamic spatiotemporal changes of ligand-receptor expression suggested that different ligand-receptor pairs are precisely programmed to coordinate the cell fate determination and organogenesis during zebrafish embryonic development (Figure 5B).

In conclusion, we demonstrated a spatiotemporal landscape of the transcriptional dynamics in the developing zebrafish embryo and provided a useful resource for studying the cellular and molecular mechanisms of germ layer specification, organogenesis and cell fate determination. Future perspectives can be the construction of the spatiotemporal zebrafish developmental atlas at a longer time window with shorter intervals, 3D reconstruction based on multiple sectioning strategy, and further exploration of the regulatory networks by integrating more spatial omics data such as the spatial genomics and chromatin accessibility.

## METHODS

### Tissue collection and sample mounting for Stereo-seq

All relevant procedures involving animal experiments presented in this paper were approved by Animal Care and Use Committee of Huazhong Agriculture University (HZAUMO-2015-016). Zebrafish embryos from AB wild-type crosses were collected at 3.3 hours, 5.25 hours, 10 hours, 12 hours, 18 hours, and 24 hours after fertilization. After being dechorionated manually with forceps, embryos were anesthetized by soaking in tricaine solution (Bomeibio, 886-86-2). Embryos were then placed in cryomold with OCT (Sakura, 4583), and extra egg water was removed using a glass pipette. The embryos were oriented with a blunt metal needle to the right position. After the OCT freezes solidly on the flat surface of a block of dry ice, the embryos were sectioned on a cryostat machine or stored at -80°C.

### Stereo-seq library preparation and sequencing

Stereo-seq library preparation and sequencing were adapted according to the standard protocol V1.1 with minor modification (Chen et al., 2021). Tissue sections were adhered to the Stereo-seq chip, and incubated in -20°C methanol for 30 min fixation, followed by nucleic acid dye staining (Thermo fisher, Q10212) and imaging (Ti-7 Nikon Eclipse microscope). For permeabilization, tissue sections before 18 hpf were permeabilized at 37°C for 3 minutes, while tissue sections of 18 hpf and 24 hpf were permeabilized at 37°C for 5 minutes. The cDNA was purified using AMPure XP beads (Vazyme, N411-03). The indexed single-cell RNA-seq libraries were constructed according to the manufacturer’s protocol. The sequencing libraries were quantified by Qubit™ ssDNA Assay Kit (Thermo Fisher Scientific, Q10212). DNA nanoballs (DNBs) were loaded into the patterned Nano arrays and sequenced on MGI DNBSEQ-Tx sequencer (50 bp for read 1, 100 bp for read 2).

### Zebrafish embryo collection and single cell isolation for scRNA-seq

The protocol for zebrafish embryos dissociation and single cell suspension isolation was adapted according to published protocol with a few modifications (Manoli and Driever, 2012). Briefly, 250–1000 embryos were collected and transferred into a Petridish containing Pronase E protease solution (Sigma-Aldrich, P5147-1G). When 20–30% embryos were hatched, 1 mL 56°C pre-heated Hi-Fetal Bovine Serum (Biological Industries, 04-001-1ACS) was added. Then embryos were washed once with 0.5X Danieau’s solution containing 10% Hi-FBS, and thrice with 0.5X Danieau’s solution. Then deyolking buffer was added and pipetted up and down until only the bodies of the embryos were visible. After dissociation with 1× trypsin-EDTA solution (Biosharp, BL512A), incubation was stopped by adding Hi-FBS to a final concentration of 5%. Finally, single cells were resuspended in 0.04% BSA (Sigma, A8022-50G) in PBS (Meilunbio, MA0015).

### Single cell RNA-seq library construction and sequencing

The DNBelab C Series Single-Cell Library Prep Set (MGI, 1000021082) was utilized according to the manufacture’s protocol (Liu et al., 2019). In brief, single cell suspensions were loaded into the chip for droplet generation. Then the droplets were gently removed from the collection and incubated at room temperature for 20 minutes to capture mRNA release from cells. And then emulsion breakage and beads collection were performed. After reverse transcription, and cDNA amplification, purified PCR products were used for DNB generation and finally sequenced on an MGI DNBSEQ-Tx using the following read length: 41bp for read1, 100bp for read2, and 10bp for sample index.

### Binning data of spatial Stereo-seq data

Raw data were processed in the same procedure as previous work (Chen et al., 2021). Transcripts captured by 15*15 DNBs were merged as one bin 15. We treated the bin 15 as the fundamental analysis unit. Bin IDs were synthesized by their spatial coordination (spatial_x and spatial_y) at the capture chip. In specific, the DNB at the left bottom of bin 15 was selected to represent the location of bin 15. Sample contours were manually drawn to exclude bins that were not from the tissue samples. For comparison with other resolutions, we also binned DNBs in other bin sizes, *i*.*e*., bin 50, 100, 150, 200, and performed unsupervised clustering. The thresholds of gene number used for filtering low-quality bins were: 150 for bin 15, 1500 for bin 50, 2500 for bin 100, 4000 for bin 150 and 5000 for bin 200.

For a demonstration of the spatial location of sparsely expressed genes in Figure 3G, we plotted every captured mRNA molecule in embryo in the resolution of bin 1 using function ‘ggplot’ in R package ggplot2 (Villanueva and Chen, 2019). And the location of molecules captured by the same DNB was jittered to avoid overlapping by using the function ‘geom_jitter’.

### Unsupervised clustering of Stereo-seq data

Data normalization, scaling, and bins clustering were processed using the R package Seurat (v3.5.1) (https://github.com/satijalab/seurat). Sections from the same embryo were pooled together as a data set for analysis. Cell identities of clusters were annotated using marker genes. Gene number captured was used for quality control, with 100 genes for 5.25 hpf and 10 hpf, 150 genes for embryos at other stages, and bins with a lower number of genes captured were filtered out. Bins located in the yolk with low gene numbers were also filtered out in this procedure. To get more detailed cell types, bins belong to a specific cell type, or groups of relevant cell types were further clustered and annotated.

### Single-cell RNA-seq data processing: cell clustering and identification of cell types

Raw sequencing reads of each sample were processed using DNBelab_C_Series_HT_scRNA-analysis-software (https://github.com/MGI-tech-bioinformatics/DNBelab_C_Series_HT_scRNA-analysis-software) which include alignment, primary filtering, and gene expression generation of each cell. We merged all data sets for each time point using the merge function and used Seurat (v4.0.3) to filter the low-quality cells by gene number of each cell and reads mapped to mitochondrial genes. The data were then log normalized using ‘NormalizeData’ function with the scaled factor set to the default value of 10,000. We scale the data using ‘ScaleData’ function and identified the 2,000 high variable features using ‘FindVariableFeatures’ function. All these variable genes were used to do principal-component analysis (PCA). The cells were clustered using ‘FindNeighbors’ function using the first 20 principal components, followed by ‘FindClusters’ function. All cell clusters were identified by using known cell type–specific markers for each cluster.

### Marker genes of clusters and GO an KEGG gene enrichment analysis

Marker genes of different clusters were found by ‘FindMarkers’ of Seurat R package, with parameter (min.pct = 0.25, logfc.threshold = 0.25). Gene set enrichment analysis, were processed with function ‘enrichGO’ using parameters (OrgDb=org.Dr.eg.db, pAdjustMethod=“BH”, pvalueCutoff = 0.01, qvalueCutoff = 0.05) and function ‘enrichKEGG’ with parameter (organism=‘dre’, pvalueCutoff=0.05) of R package clusterProfiler (https://bioconductor.org/packages/release/bioc/html/clusterProfiler.html).

### Identification of spatially auto-correlated gene modules

Modules of spatially correlated genes were identified using Hotspot (DeTomaso and Yosef, 2021). The expression matrixes of genes with minimum UMIs number (50 for 10, 12, and 18 hpf; 300 for 24 hpf) of all embryo sections from 4 development stages (10, 12, 18 and 24 hpf) were used as input. The data were first normalized by the total UMIs number of these genes of each bin, then K-Nearest Neighbors (KNN) graph of genes was created using the ‘create_knn_graph’ function with the parameters: n_neighbors = 8, then genes with significant spatial autocorrelation (FDR < 0.05) were kept for further analysis. The modules were identified using the ‘create_modules’ with the parameters: min_gene_threshold (20 genes for 10 and 12 hpf, 30 genes for 18 hpf, and 50 genes for 24 hpf) and fdr_threshold = 0.05. For the cell type enrichment analysis for each module, we calculated the cell type composition of bins with high module score (with the criteria that module scores bigger than 3, the method of calculation of the module score see the Hotspot reference (DeTomaso and Yosef, 2021)). As a comparison, we also calculated the cell type composition of all sections of the same developmental stage (Figure 3B, S2B, S2E and S2H).

To find genes that spatially correlated with Notch pathway, we further conducted module analysis in selected sections at 12, 18 and 24 hpf (section #8, #8, and #4, with minimum UMIs number for gene selection as 50, 30, 50 respectively.). We first calculated gene set expression score (Notch-score) of Notch pathway genes (genes with very low expression were removed from the Notch pathway gene list) using function ‘AddModuleScore’ of R package Seurat. Then we added the Notch-score into the gene expression matrix and calculated the spatial correlations as above. Genes with Z-scores for the significance of the correlation with Notch-score bigger than 5 were chosen for further analysis. Modules of these Notch pathway correlated genes were identified using ‘create_modules’, with parameter min_gene_threshold, 5, 10, 20 for 12, 18, 24 hpf respectively. Notch genes used for analysis included: *dlc, dla, dld, dlb, dll4, jag1a, jag1b, jag2b, notch1a, notch1b, notch2, notch3, her6, her9, her12, her15*.*1, her15*.*2, her4*.*2, her2, her4*.*4, her3, her13, her8*.*2, her8a, her4*.*1, her4*.*3, her4*.*5, her7, hey1, hey2*.

### Protein-protein interaction (PPI) network analysis of spatial modules

We applied the search tool for the retrieval of interacting genes (STRING) (https://string-db.org) database to seek potential interactions among genes in one/multiple modules. Active interaction sources including co-expression as well as species limited to “*Danio rerio*”, and an interaction score >0.4 were applied to construct the PPI networks. The PPI networks were visualized by Cytoscape software 3.8.2. In these networks, the nodes correspond to the proteins and the node size is proportional to the relative connectivity in each network. The dark lines represent positive associations and the thickness represents the strength of the interactions. We chose “co-expression” as edge weights to represent the strength of the interactions.

### Construction of single-cell developmental trajectory

We integrated scRNA-seq data from adjacent time points using Harmony (lambda = 0.1, epsilon.harmony = -Inf) (Korsunsky et al., 2019). Then we calculated the 10 nearest neighbors in the latter time point for each spot of the earlier time point using K-Nearest Neighbors algorithm. We took the most frequent clusters of the 10 nearest spots as the target states that the earlier spots would develop into. If two or more target clusters had the same frequencies in the nearest neighbors, the cluster with the shortest weighted distance was used. We repeated this procedure on each pair of adjacent time points from 3.3 hpf to 24 hpf. While constructing the developmental trajectory between each two adjacent time points, we reserved only the connections with a proportion of source cells more than 30% of its type.

### Deconvolution of cell types

We divided single-cell clusters into subtypes according to the single-cell developmental trajectory. Then we calculated the cell composition of each spatial data bin using SPOTlight (Elosua-Bayes et al., 2021) with scRNA-seq data as a reference.

### TF regulation activity prediction

We downloaded the motif database of zebrafish from the CIS-BP Database (http://cisbp.ccbr.utoronto.ca/bulk.php) and constructed the cisTarget databases for zebrafish according to the SCENIC tutorial instructions (Aibar et al., 2017) (https://github.com/aertslab/create_cisTarget_databases). Then we calculated the single-cell TF regulation activity using pySCENIC (rank_threshold = 10000, auc_threshold = 0.1, nes_threshold = 2.5). We used the weighted average expression of predicted target genes to illustrate the regulation activity of each transcription factor in the spatial transcriptomic data.

### Ligand-receptor analysis

Ligand-receptor pairs were extracted from the LRBase.Dre.eg.db database (http://bioconductor.org/packages/release/data/annotation/html/LRBase.Dre.eg.db.html). For each bin expressing a receptor, we calculated its Euclidian distances to all the bins expressing the corresponding ligand and take the nearest distance as the distance of this ligand-receptor pair in this bin. Then the distances were scaled regarding the distance of adjacent cells as 1. The average distance of each ligand-receptor pair in each time point was taken as the ligand-receptor distance for that time point. Ligand-receptor pairs with an average distance less than 5 in one or more time points were kept for analysis. The distance score was calculated as:

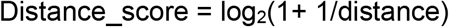

We calculated the expression score of each ligand-receptor pair as:

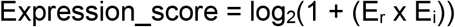

Where E_r_ and E_i_ represent the average expression of the receptor and the ligand respectively.

For receptor-ligand interaction of notch signal, we normalized the interaction frequency by dividing the bin number of each cell type.

## ACKNOWLEDGEMENTS

This work was supported by National Natural Science Foundation of China (Grant No. 31871481, 32070973, 31601181, 31571107). We thank Dr. A.F. Schier for his valuable suggestions on data analysis and for critically reading the manuscript.

## AUTHOR CONTRIBUTIONS

X.X., Z.D. and L.L. designed and supervised the work. C.L., X.X., Z.D. and L.L. designed the experiment. C.L., X.L., S.W., K.Z., X.S., C.P., H.W. and W.B. performed the library preparation and sequencing. R.L., Young.L, Q.L. Y.M. and X.Y. performed the bioinformatics analysis. K.M. and J.H. provided technical support. A.C. and Yuxiang.L. gave the relevant advice. M.A.E., M.A.B., S.K.S., S.L. and X.X. participated in the manuscript editing and discussion. C.L., Z.D. and L.L. wrote the manuscript. All authors edited and approved the manuscript.

## DATA AND CODE AVAILABILITY

All raw data have been deposited to CNGB Nucleotide Sequence Archive (accession code: CNP0002220).

## FIGURE LEGENDS

**Figure S1.**
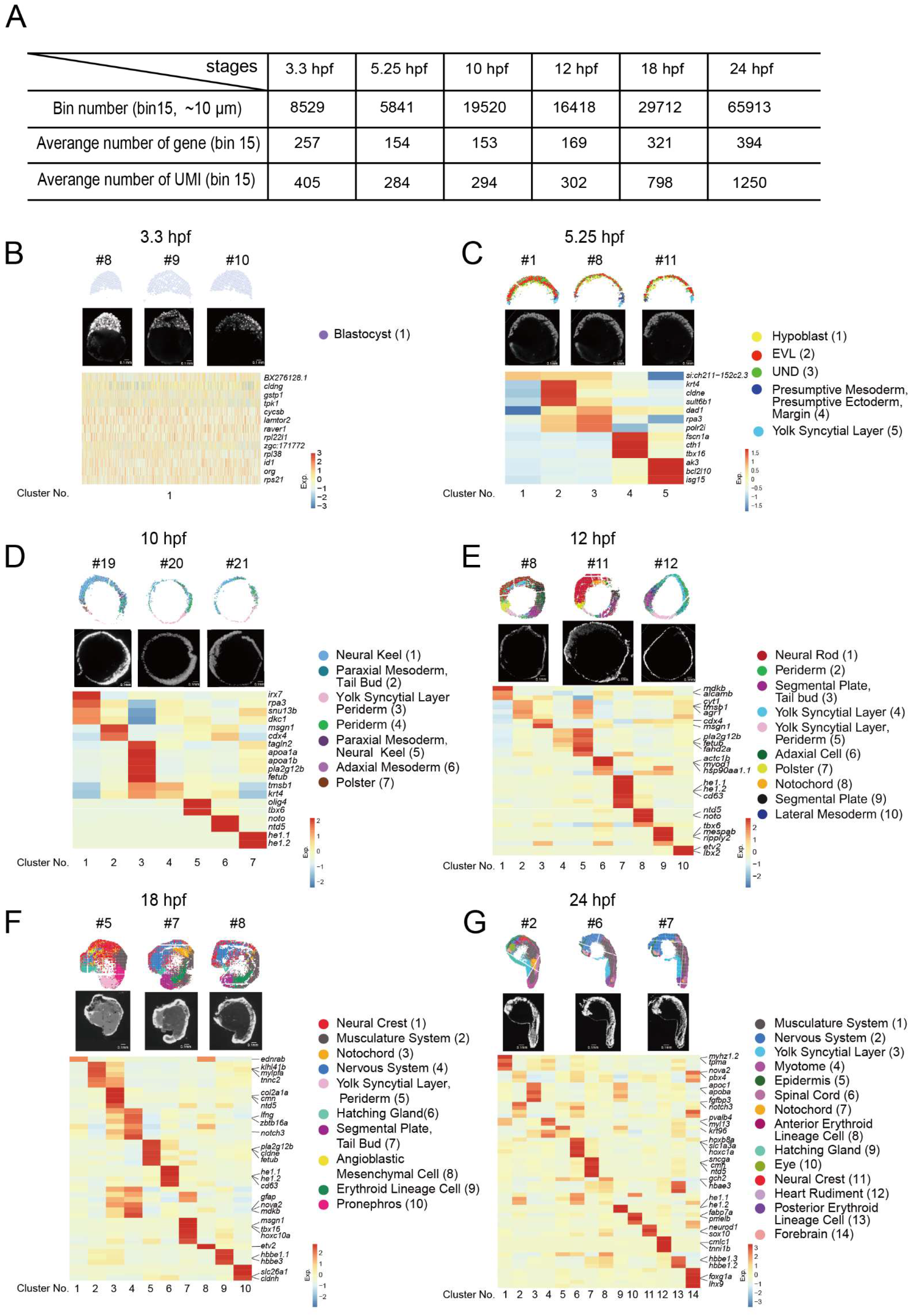
Spatially resolved transcriptomic atlas across 6 sequential developmental stages. Related to Figure 1 and 2. (A) The summary of total bins, average captured number of UMIs and genes at bin 15 resolution for each embryonic development stage. (B-G) Unsupervised clustering of 3 selected sections from 3.3 hpf (B), 5.25 hpf (C), 10 hpf (D), 12 hpf (E), 18 hpf (F) and 24 hpf (G) embryos by Stereo-seq (top) and the images of nucleic acid dye staining of the same embryo sections (middle). Scale bars: 0.1 mm. Heatmap shows the expression pattern of marker genes of different clusters at each developmental stage (bottom).

**Figure S2.**
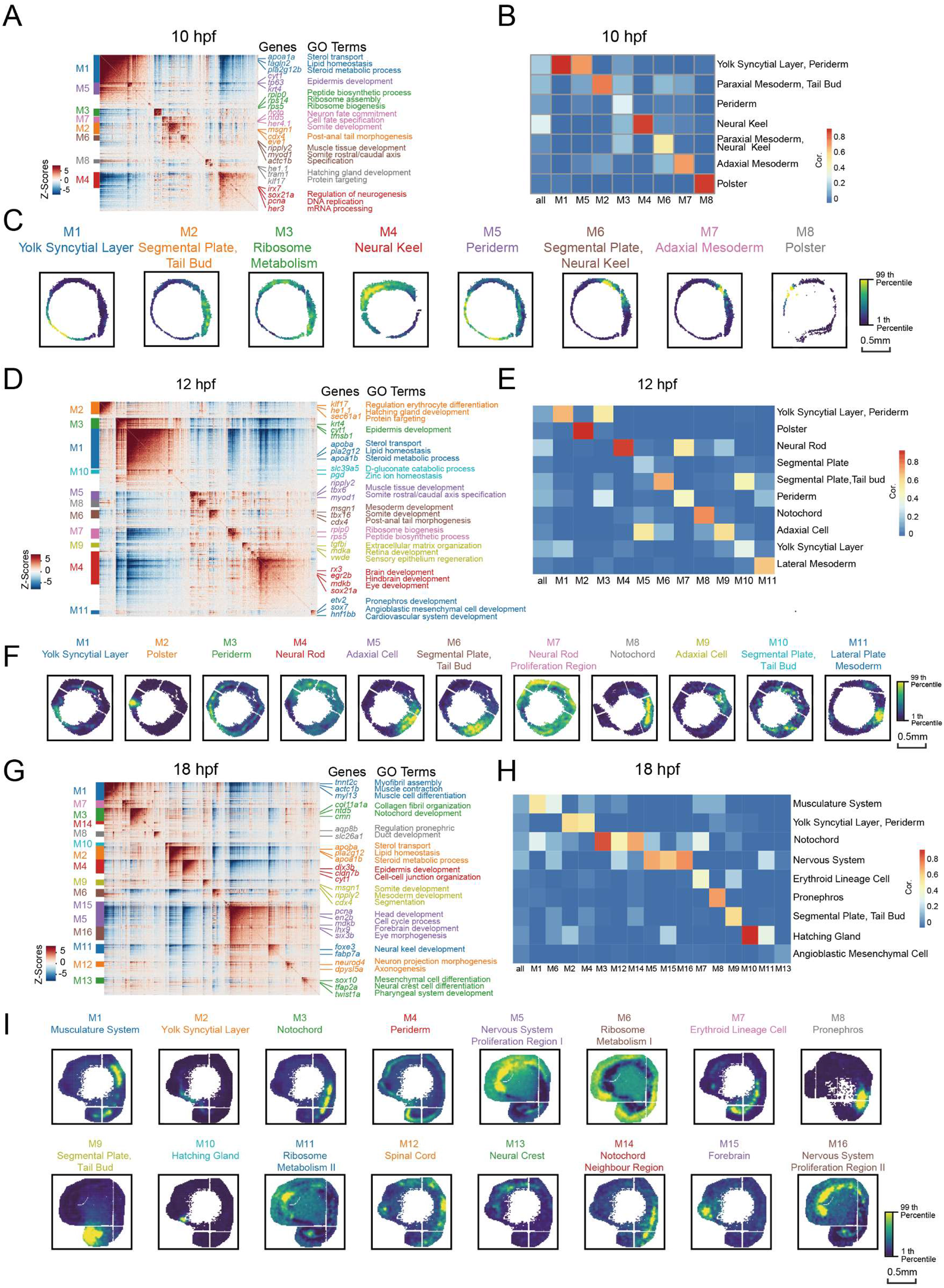
Hotspot identified spatial modules at multiple zebrafish embryo sections at different developmental stages. Related to Figure 3. (A, D, G) Heatmap shows the genes with significant spatial autocorrelation (449 genes at 10 hpf, 791 genes at 12 hpf, 2320 genes at 18 hpf, FDR<0.05) grouped into 8 gene modules at 10 hpf (A), 11 gene modules at 12 hpf (D) and 16 gene modules at 18 hpf (G). Selected genes and representative GO terms related to the corresponding modules are highlighted on the right side. (B, E, H) Heatmap shows the Pearson correlation of the module score for each spatial autocorrelation module and the expression sets of specific genes for each spatial cluster of 10 hpf (B), 12 hpf (E) and 18 hpf (H) embryos. (C, F, I) Spatial visualization of modules on 10 hpf (C), 12 hpf (F) and 18 hpf (I) embryo sections. Scale bars: 0.5 mm.

**Figure S3.**
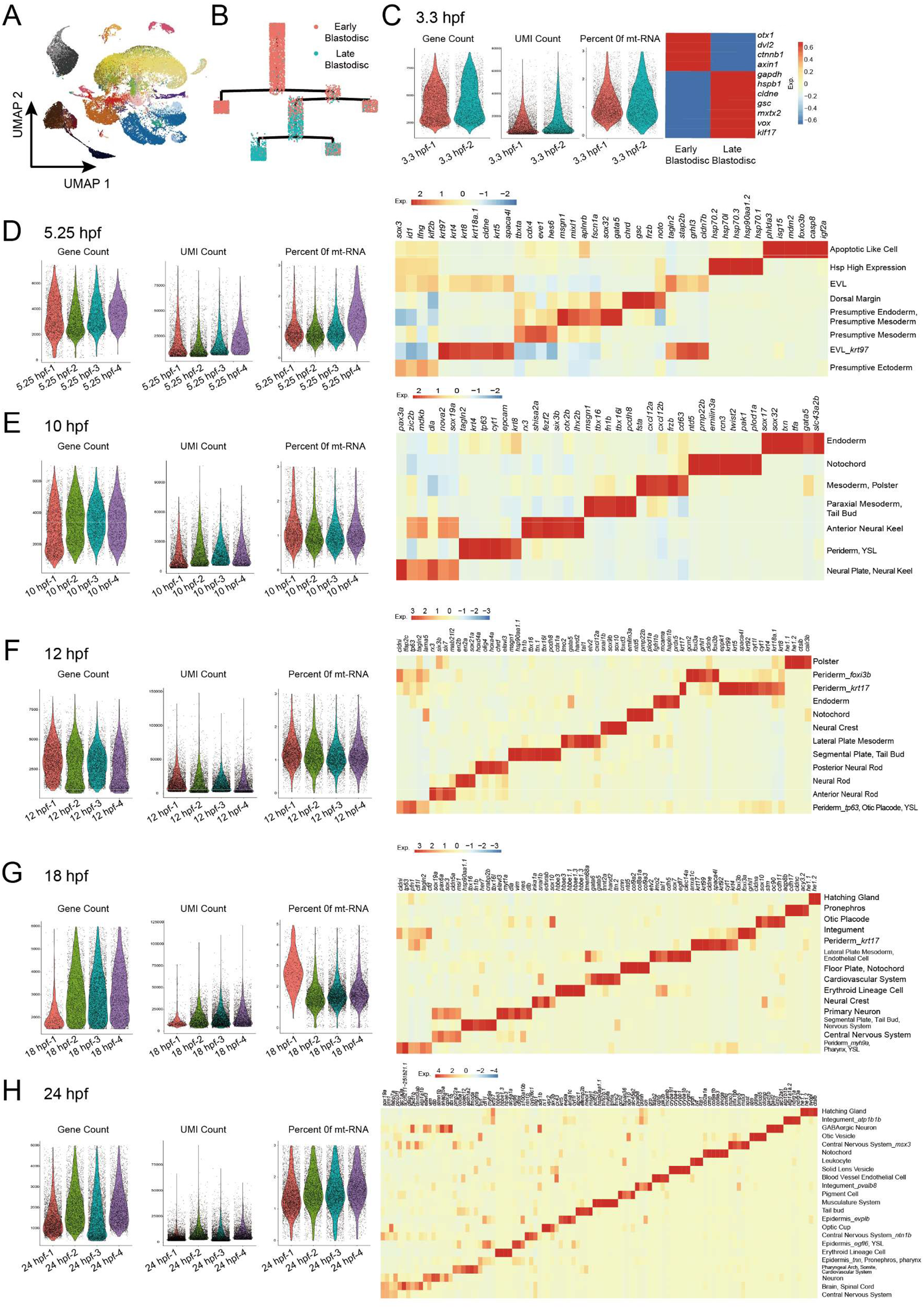
Quality control of the single-cell RNA-seq libraries. Related to Figure 4. (A) UMAP visualization of all single-cell RNA-seq data of the 6 developmental stages. Visualization of each cell type colored by tissue/organ. The color legend of A is the same as that in Figure 4B. (B) Pseudotime analysis of the 3.3 hpf embryo is taken by Monocle 2, and cells are colored by cell type. (C-H) Violin plots show the number of genes (left), the number of UMIs (middle left), and the percentage of mitochondrial genes (middle right) from multiple library repetitions with 3.3 hpf (C), 5.25 hpf (D), 10 hpf (E), 12 hpf (F), 18 hpf (G), and 24 hpf (H) zebrafish embryos. Heatmaps showing expression of marker genes of the indicated cell type of each development stage (right).

**Figure S4.**
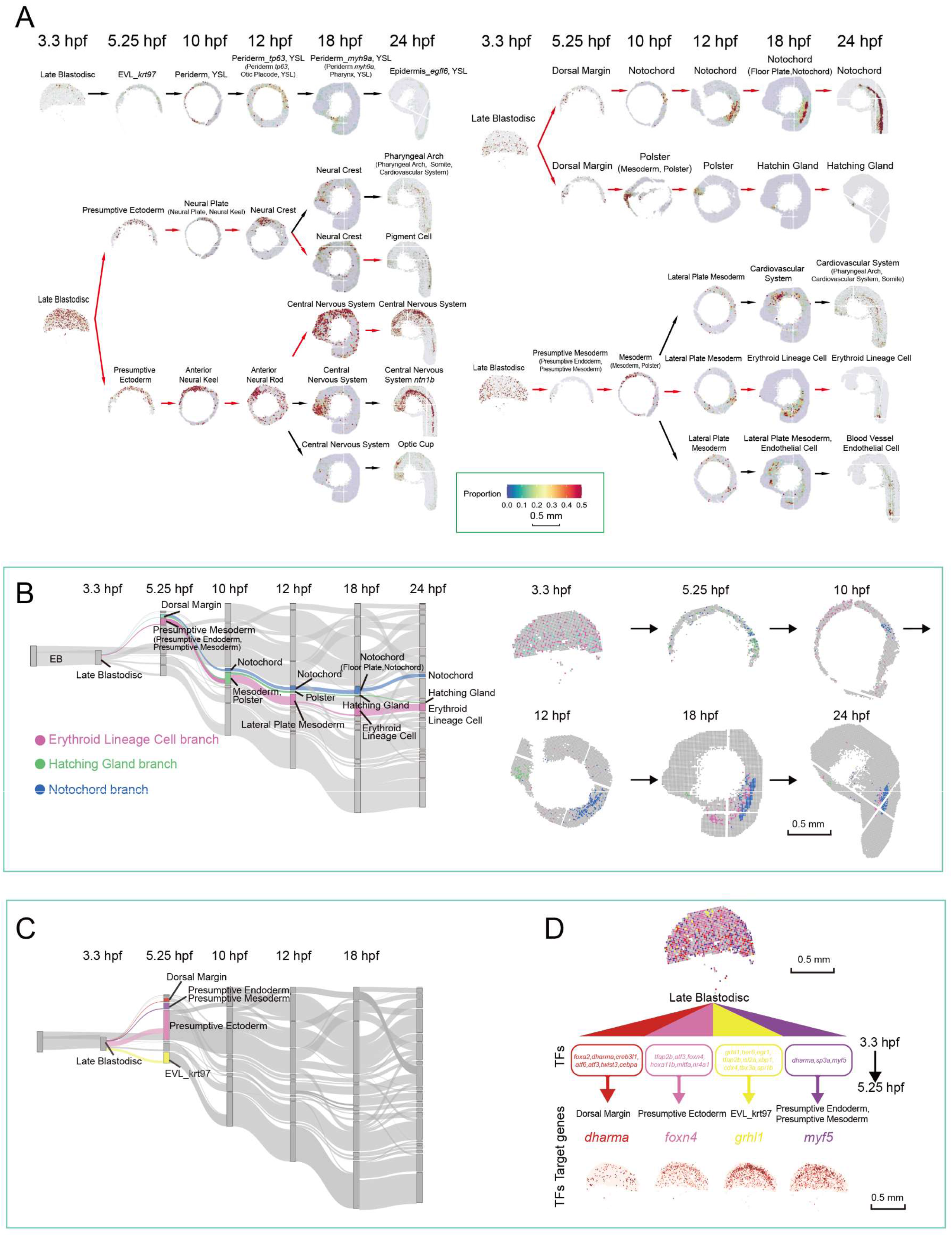
The selected spatially resolved developmental trajectories. Related to Figure 4. (A) The representative developmental branches selected from Figure 4B are displayed respectively on sequential spatial sections across all developmental time points to show the spatial trajectory. The red arrows indicated the branches shown in Figure 4C and S4B. (B) Application of SPOTlight to integrate Stereo-seq with scRNA-seq data to infer the spatially resolved developmental trajectories. Three developmental branches which are developed from the dorsal margin and presumptive mesoderm, namely, erythroid lineage cell, hatching gland and notochord are simultaneously displayed on spatial sections. (C) The selected developmental branching points at 3.3 hpf are shown in the Sankey diagram. (D) Cell fate regulatory maps of different developmental destinations at 3.3 hpf. Scale bar: 0.5 mm. SPOTlight was applied to visualize the spatial location of cell subgroups with different differentiation (top); crucial TFs for different developmental branches are shown in the chart (middle); the spatial expression distributions of TFs target genes are visualized on embryonic sections (bottom).

**Figure S5.**
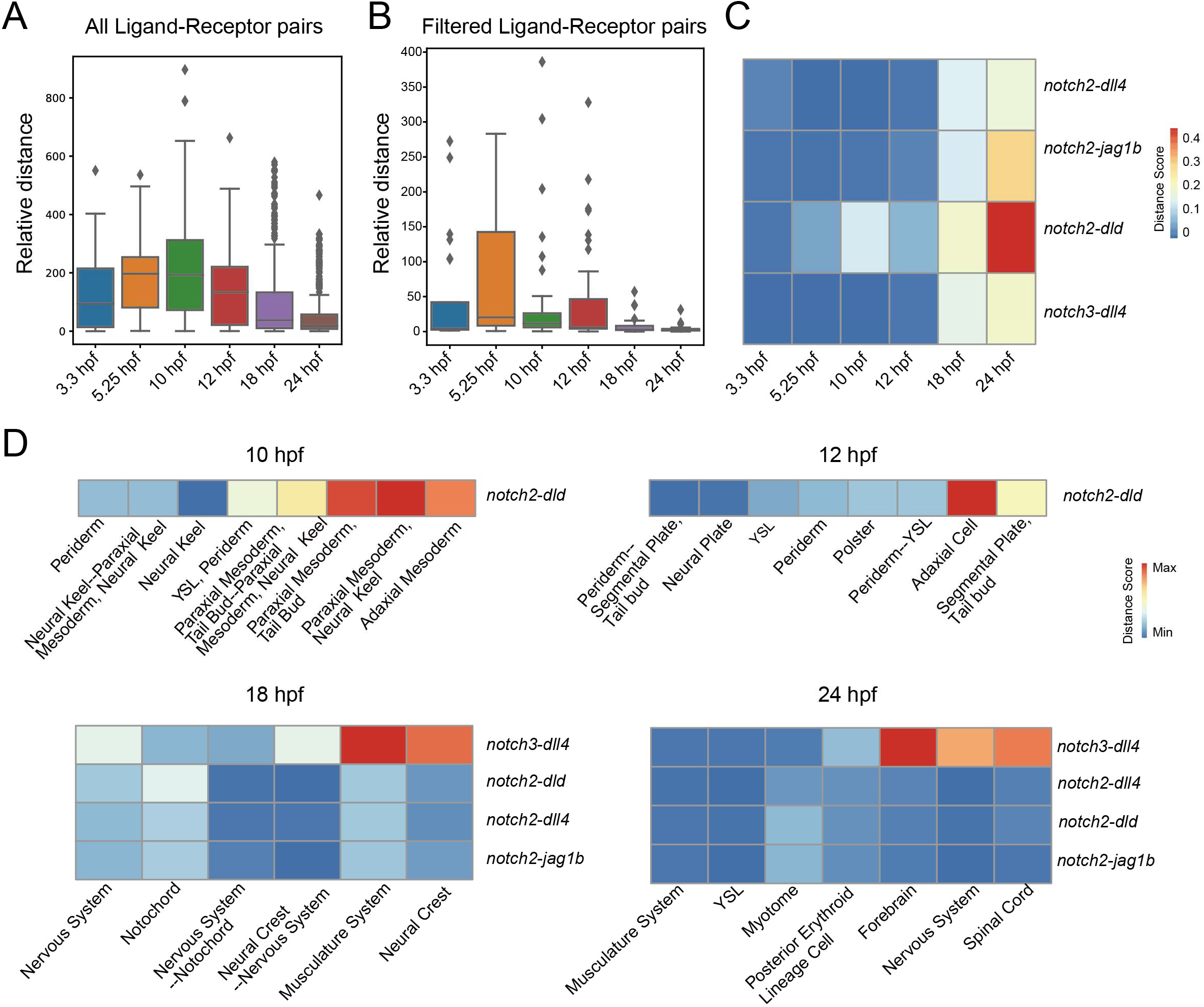
Analysis of the relative spatial distance of ligand-receptor pairs during zebrafish embryo development. Related to Figure 5. (A) Boxplot shows the relative distance values of all ligand-receptor pairs at different time points. (B) Boxplot shows the relative distance values of filtered ligand-receptor pairs at the different time points. Filtered ligand-receptor pairs refer to those which have a relative distance less than 5 for at least one time point. (C) Heat map shows the relative distance of Notch ligand-receptor pairs. (D) Heat maps show the relative distance between Notch ligands and receptors in representative cell types at 10 hpf, 12 hpf, 18 hpf and 24 hpf.

